# Transcriptome-wide association studies: opportunities and challenges

**DOI:** 10.1101/206961

**Authors:** Michael Wainberg, Nasa Sinnott-Armstrong, Nicholas Mancuso, Alvaro N. Barbeira, David A Knowles, David Golan, Raili Ermel, Arno Ruusalepp, Thomas Quertermous, Ke Hao, Johan LM Björkegren, Hae Kyung Im, Bogdan Pasaniuc, Manuel A Rivas, Anshul Kundaje

## Abstract

Transcriptome-wide association studies (TWAS) integrate GWAS and gene expression datasets to find gene-trait associations. In this Perspective, we explore properties of TWAS as a potential approach to prioritize causal genes, using simulations and case studies of literature-curated candidate causal genes for schizophrenia, LDL cholesterol and Crohn’s disease. We explore risk loci where TWAS accurately prioritizes the likely causal gene, as well as loci where TWAS prioritizes multiple genes, some of which are unlikely to be causal, because they share the same variants as eQTLs. We illustrate that TWAS is especially prone to spurious prioritization when using expression data from tissues or cell types that are less related to the trait, due to substantial variation in both expression levels and eQTL strengths across cell types. Nonetheless, TWAS prioritizes candidate causal genes at GWAS loci more accurately than simple baselines based on proximity to lead GWAS variant and expression in trait-related tissue. We discuss current strategies and future opportunities for improving the performance of TWAS for causal gene prioritization. Our results showcase the strengths and limitations of using expression variation across individuals to determine causal genes at GWAS loci and provide guidelines and best practices when using TWAS to prioritize candidate causal genes.

---

Over the past 13 years, genome-wide association studies (GWAS) have robustly associated thousands of genomic loci with a variety of complex traits. Despite this success, GWAS loci are often difficult to interpret: linkage disequilibrium (LD) often obscures the causal variant(s) driving the association, and the causal genes mediating these variants’ effects on the trait can rarely be ascertained from GWAS data alone^1^. This interpretational challenge has motivated the development of methods to prioritize causal genes at GWAS loci.

One such family of methods are transcriptome-wide association studies (TWAS). TWAS leverage expression reference panels (eQTL cohorts with expression and genotype data) to discover gene-trait associations from GWAS datasets^2,3,4^. First, the expression reference panel is used to learn predictive models of expression variation for each gene using allele counts of genetic variants in the vicinity of the gene (typically 500 kb or 1 MB around the gene). Next, these models are used to predict gene expression for each individual in the GWAS cohort. Finally, statistical associations are estimated between predicted gene expression and the status of the trait (Fig. 1). The expression prediction and association may be performed explicitly using individual-level GWAS data, as in PrediXcan^2^, or combined into a single step using summary-level GWAS data, as in Fusion^3^ and S-PrediXcan^4^. Closely related methods to TWAS include SMR/HEIDI^5,6,7^, which performs Mendelian Randomization (MR) from gene expression to trait, and GWAS-eQTL colocalization methods such as Sherlock^8^, coloc^9,10^, QTLMatch^11^, eCaviar^12^, enloc^13^ and RTC^14^, which discover genes whose expression is regulated by the same causal variant(s) that underlie a GWAS hit.

**Fig. 1:**
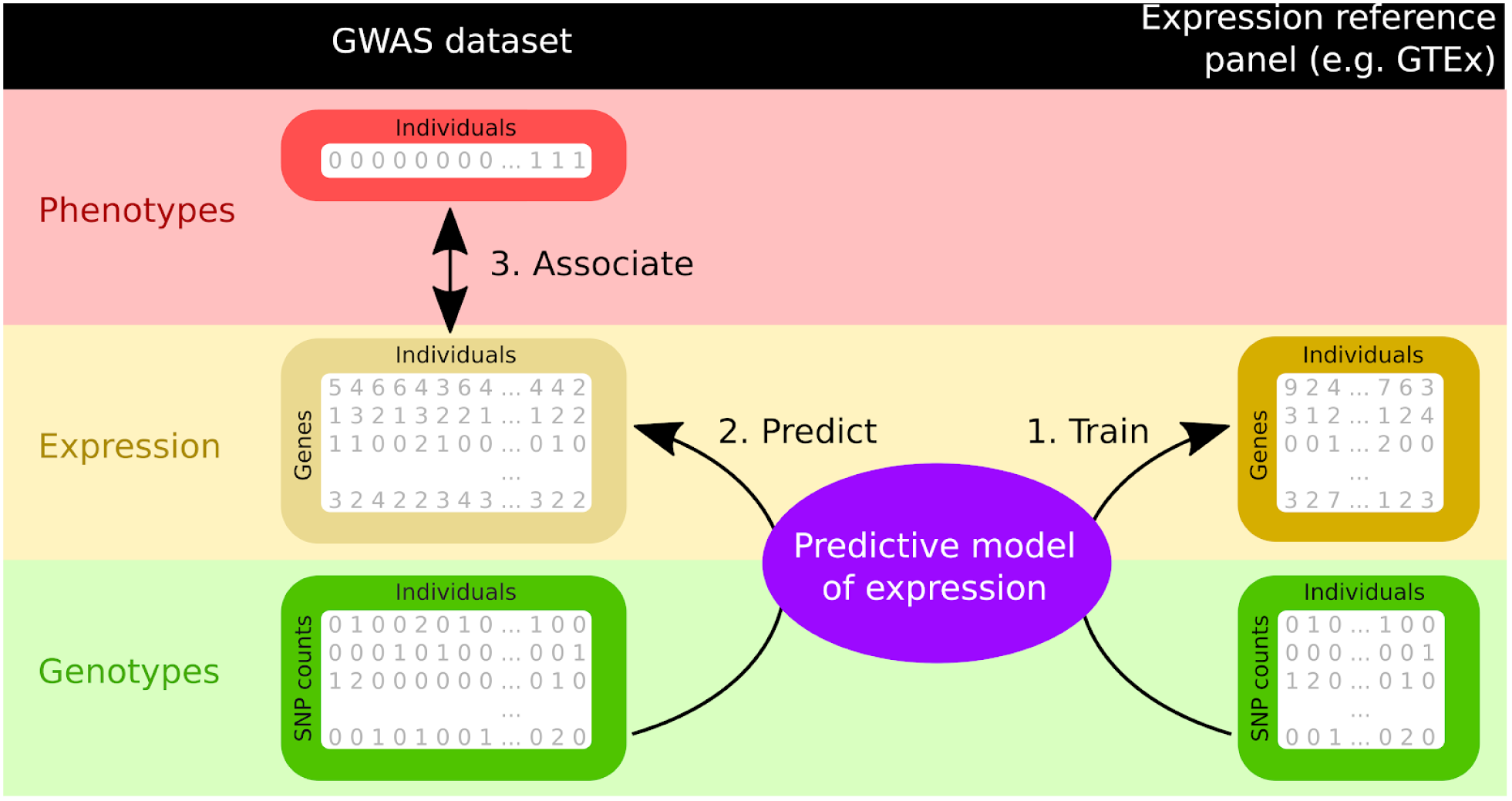
An overview of TWAS. Briefly, TWAS involves: 1) training a predictive model of expression from genotype on a reference panel such as GTEx; 2) using this model to predict expression for individuals in the GWAS cohort; and 3) associating this predicted expression with the trait.

TWAS have garnered substantial interest within the human genetics community and have subsequently been conducted for a wide variety of traits and tissues^15,16^. Although TWAS has been proposed as a statistical test for finding associations between the genetic component of expression and disease risk (with no causality guarantees), a key reason for the appeal of TWAS methods is the promise that they may prioritize candidate causal genes (defined here as genes that mediate the phenotypic effects of causal genetic variants) at known GWAS risk regions. Unfortunately, there is a prevalent misconception within the community that TWAS is a causal gene test and that TWAS associations represent *bona fide* causal genes; in the following sections, we provide guidelines for interpreting TWAS results, highlighting situations when TWAS can accurately prioritize candidate causal genes as well as scenarios when the genes prioritized by TWAS are likely non-causal.

As a motivating example illustrating both the success and interpretational challenges of TWAS, consider *C4A*, a causal gene for schizophrenia. Variants at the *C4A* locus contribute to schizophrenia risk by increasing expression of *C4A* in the brain^17^. TWAS detects a strong association between *C4A* and schizophrenia risk using brain expression data from GTEx^18^. Strikingly, *C4A* is by far the most significantly associated gene within the 100 kb locus in brain tissues. C4A is also the the most significantly associated gene in any tissue (Table 1), even compared to other closely-related genes in the complement system (*C4B*, *CFB*, *C2*). However, 8 of the 12 other genes within 100 kb are also at least marginally significant (p < 0.05) in some brain tissue, and 11 of 12 are highly significant (p < 5e-5) in at least one tissue.

**Table 1:** The *C4A* locus, a success story where TWAS p-values accurately prioritize the causal gene. Lowest schizophrenia p-value in any GTEx brain tissue, and in any GTEx tissue, for each gene within 100 kb of *C4A* with available S-PrediXcan TWAS results (http://metabeta.gene2pheno.org). The TWAS used schizophrenia summary GWAS data from the Psychiatric Genomics Consortium^19^ and expression data from GTEx^18^.

| Gene | Lowest TWAS p-value in any brain tissue | Lowest TWAS p-value in any tissue |
| --- | --- | --- |
| *C4A* | 4e-18 (Hypothalamus) | 2e-20 (Pancreas) |
| *ATF6B* | 3e-9 (Anterior cingulate cortex) | 3e-9 (Anterior cingulate cortex) |
| *CYP21A2* | 5e-7 (Cortex) | 9e-19 (Aorta) |
| *NELFE* | 7e-7 (Cerebellum) | 7e-7 (Cerebellum) |
| *STK19* | 1e-5 (Frontal Cortex, BA9) | 4e-12 (Adrenal gland) |
| *SKIV2L* | 5e-5 (Cerebellum) | 5e-5 (Cerebellum) |
| *C4B* | 6e-5 (Nucleus accumbens, basal ganglia) | 1e-21 (Testis) |
| *C2* | 0.008 (Cortex) | 1e-18 (Whole blood) |
| *DXO* | 0.03 (Putamen, basal ganglia) | 0.02 (Thyroid) |
| *CFB* | Not significant | 2e-13 (Whole blood) |
| *EHMT2* | Not significant | 3e-10 (Skin, sun-exposed lower leg) |
| *TNXB* | Not significant | 3e-6 (Adrenal gland) |
| *ZBTB12* | Not significant | 7e-5 (Ovary) |

## TWAS loci frequently contain multiple associated genes

It is well known that GWAS rarely identifies single variant-trait associations, but instead identifies blocks of associated variants in linkage disequilibrium with each other (Fig. 1a). Analogously, TWAS frequently identifies multiple hit genes per locus (Fig. 1b)^16^.

**Figure 1b:**
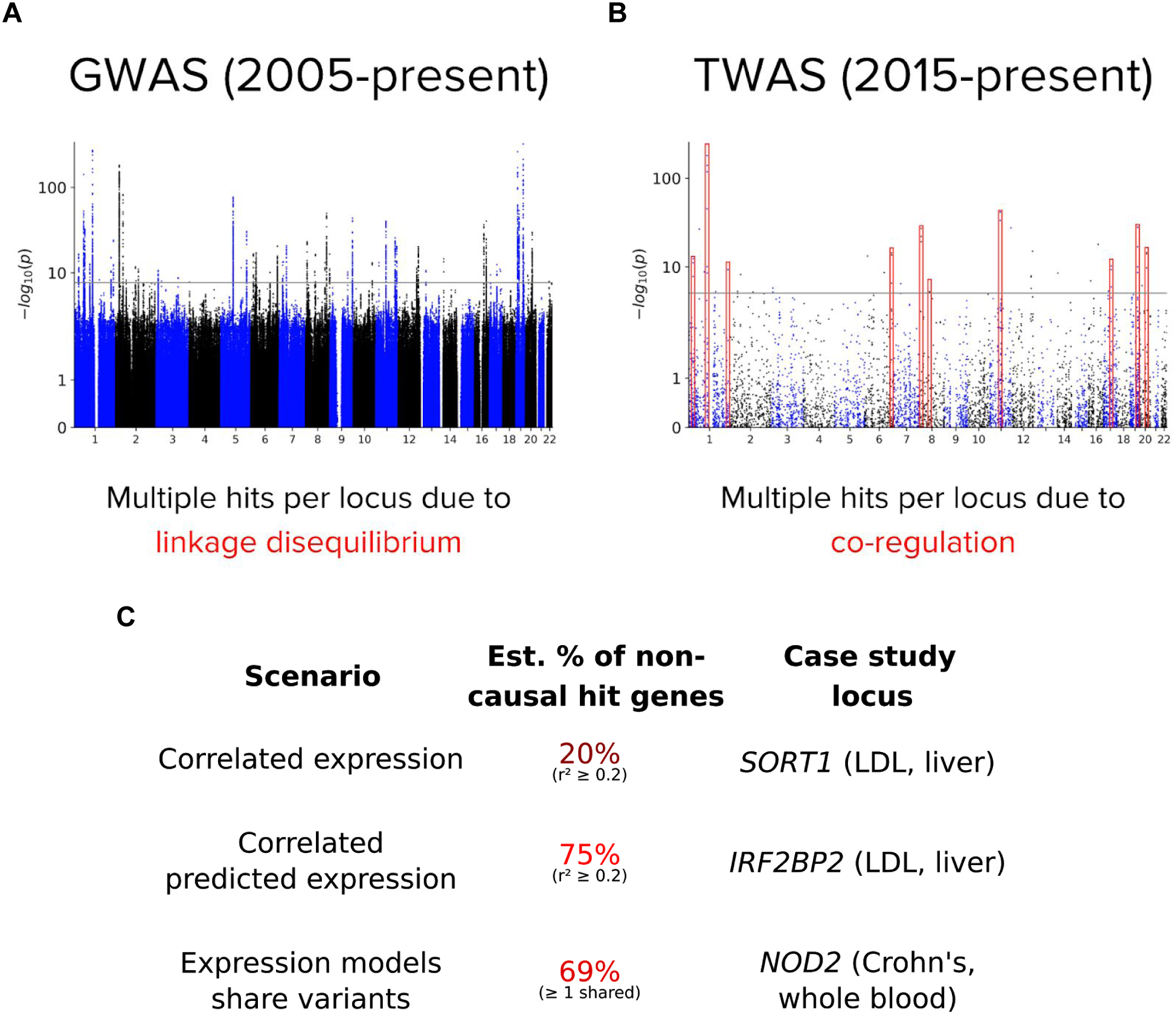
TWAS, like GWAS, frequently has multiple significant associations per risk region. (a), (b) Manhattan plots of GWAS and Fusion TWAS for LDL cholesterol using GWAS summary statistics from the Global Lipids Genetics Consortium and liver expression from the STARNET cohort (see Methods). GWAS has multiple hits per locus due to linkage disequilibrium, and TWAS due to co-regulation (which can also be driven in part by LD; see below), as we explore in the paper. Clusters of multiple adjacent TWAS hit genes are highlighted in red. (c) Three scenarios where co-regulation can lead to multiple hits per locus, and the estimated percent of non-causal hit genes subject to each scenario; each scenario is presented in a case study later in the paper (a fourth scenario is presented in Fig. 5d). To estimate the percentages, we group hits into 2.5 MB clumps and make the approximation that genes that are not the top hit in multi-hit clumps are non-causal; we then calculate the percent of these genes with total or predicted expression r^2^ ≥ 0.2 or ≥ 1 shared variant with the top hit in their block, aggregating genes across the LDL/liver and Crohn’s/whole blood TWAS. The full distributions of total and predicted expression correlations and number of shared variants are shown in Fig. S1, separated by study.

To explore this phenomenon, we performed TWAS in two traits and two tissues with both Fusion and S-PrediXcan, using GWAS summary statistics for LDL cholesterol^20^ and Crohn’s disease^21^ and the 522 liver and 447 whole blood expression samples from the STARNET cohort^22^ (Fig. S2, Online Methods). We grouped hit genes within 2.5 MB and found that while some loci contained only a single hit gene, many contained two, three, four or even up to eleven (Fig. S3).

## Correlated expression across individuals may lead to spurious TWAS hit genes

We explored the extent to which co-regulation could be responsible for multi-hit loci. The conventional way co-regulation is measured is by correlating the expression of a pair of genes across individuals in an expression cohort. Do genes that have correlated expression with a strong TWAS hit also tend to be TWAS hits (Fig. 5a)? We analyzed the *SORT1* locus in LDL/Liver (TWAS p < 1 × 10^−243^; Fig. 2a) which represents the strongest hit locus across all four Fusion TWAS. *SORT1* has strong evidence of causality (Table 3).

**Figure 2:**
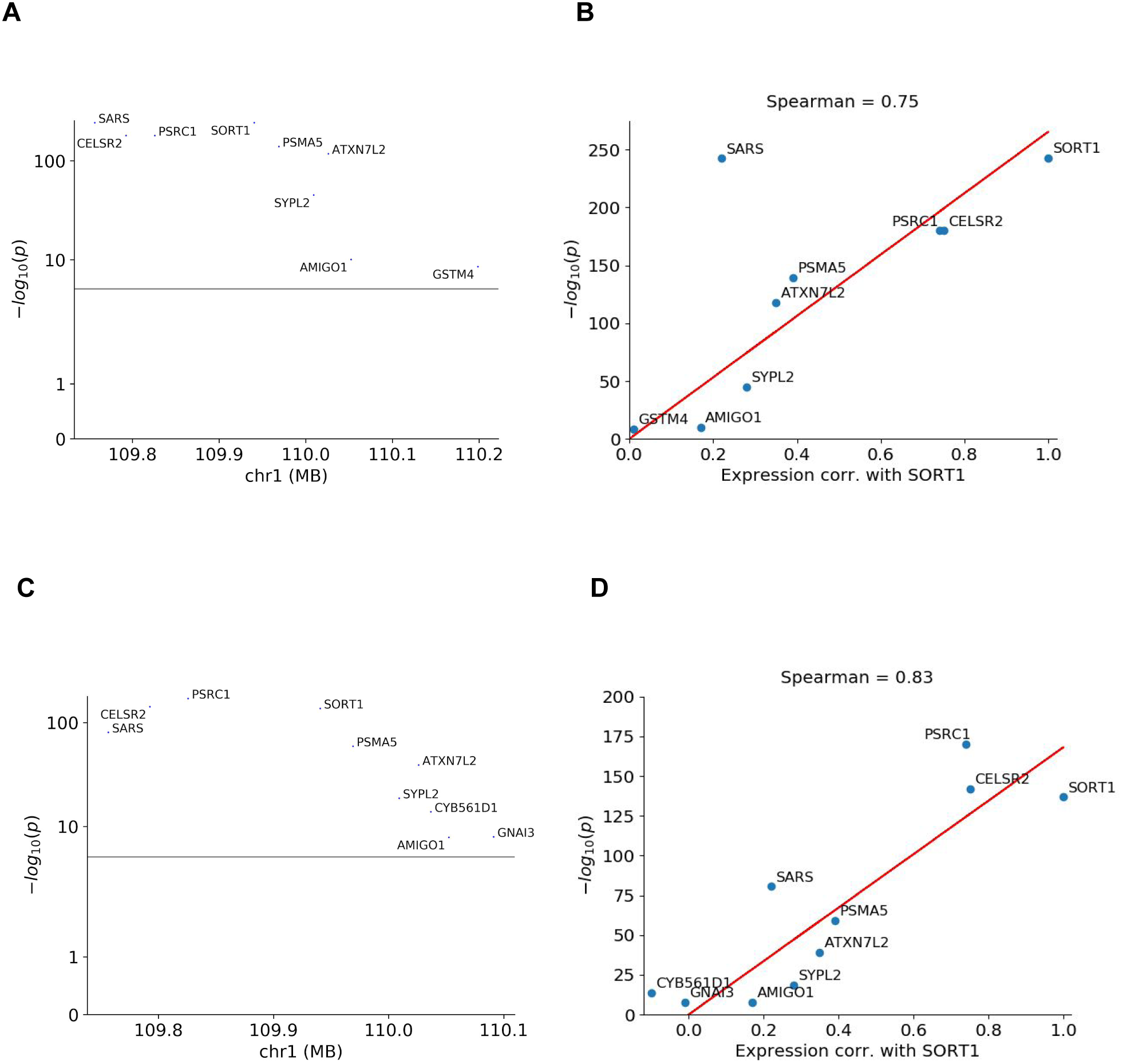
Co-regulation strongly predicts TWAS hit strength at the *SORT1* locus. a) Fusion Manhattan plot of the *SORT1* locus. b) Expression correlation with *SORT1* versus TWAS *p*-value, for each gene in the *SORT1* locus. c), d) The equivalent Manhattan and expression correlation plots for S-PrediXcan.

The *SORT1* locus contains 8 other Fusion hit genes besides *SORT1*, and their TWAS *p*-values are highly related to their expression correlation with *SORT1* (Spearman = 0.75; Fig. 2b). A similar pattern holds for S-PrediXcan (Fig. 2c, d). The two genes with the highest expression correlations with *SORT1*, *PSRC1* and *CELSR2*, were previously noted by one of the *SORT1* mouse model studies^23^ to share an eQTL with *SORT1* in liver (rs646776). Given that *SORT1* has strong evidence of causality, and that other genes at the locus lack strong literature evidence, the most parsimonious explanation is that most or all of the other genes are non-causal and are only prioritized due to their correlation with *SORT1*. However, we emphasize that there is no guarantee that other genes at the locus are truly non-causal, and some or all of these genes could be causal to various degrees.

**Table 2:**
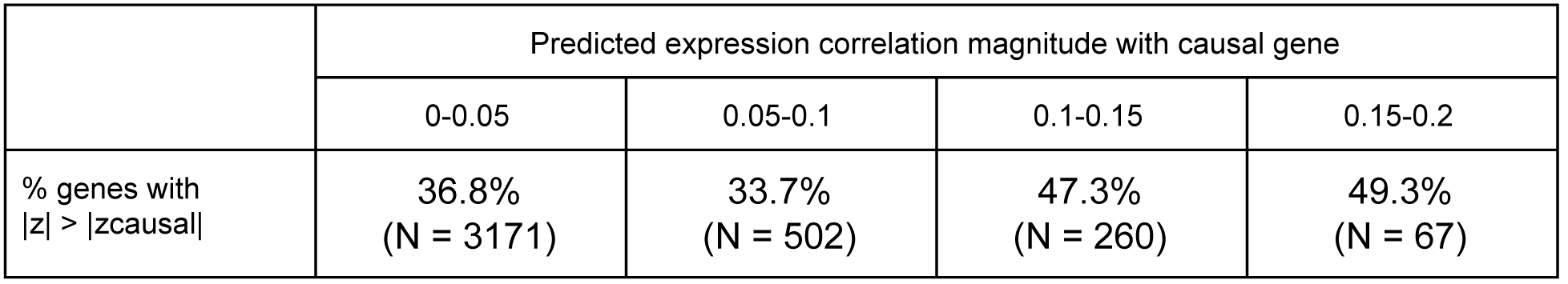
Simulation of percent of genes with larger TWAS z-score than the causal gene, binned by predicted expression correlation. The number of genes in each bin (among all genes at the 1000 random loci being simulated) is shown in brackets for each bin. Predicted expression correlations were computed as the vector-matrix-vector product of the causal gene’s model weights, the LD matrix among the variants included in the models, and the other gene’s model weights.

## Correlated predicted expression across individuals may lead to spurious TWAS hit genes, even without correlated total expression

However, expression correlation is not the whole story: after all, TWAS tests for association with genetically-predicted expression, not total expression. Total expression includes genetic, environmental and technical components, and the genetic component of expression includes contributions from common *cis* eQTLs (the only component reliably detectable in current TWAS methods), rare *cis* eQTLs, and *trans* eQTLs. Predicted expression likely only represents a small component of the GWAS individuals’ total expression: a large-scale twin study^24^ found that common *cis* eQTLs explain only about 10% of genetic variance in gene expression.

While predicted expression correlations between genes at the same locus are often similar to total expression correlations, they are generally slightly higher, and sometimes substantially so (Fig. 3a, Fig. 3d, Fig. S4). A pair of genes can have correlated predicted expression if a) the same causal eQTL regulates both genes, or b) two causal eQTLs in LD each regulate one of the genes^25^. Although only case a) counts as mechanistic co-regulation, we consider both cases together since they are not designed to be distinguishable by TWAS: the two genes’ TWAS models can rely on distinct variants even in case a), or rely on the same variant even in case b). For instance, when faced with a causal eQTL in near-perfect LD with another variant, an L1-penalized linear expression model (lasso) will generally include only one of the two variants, but which variant is chosen could flip based on stochastic statistical fluctuations in the training set.

**Figure 3:**
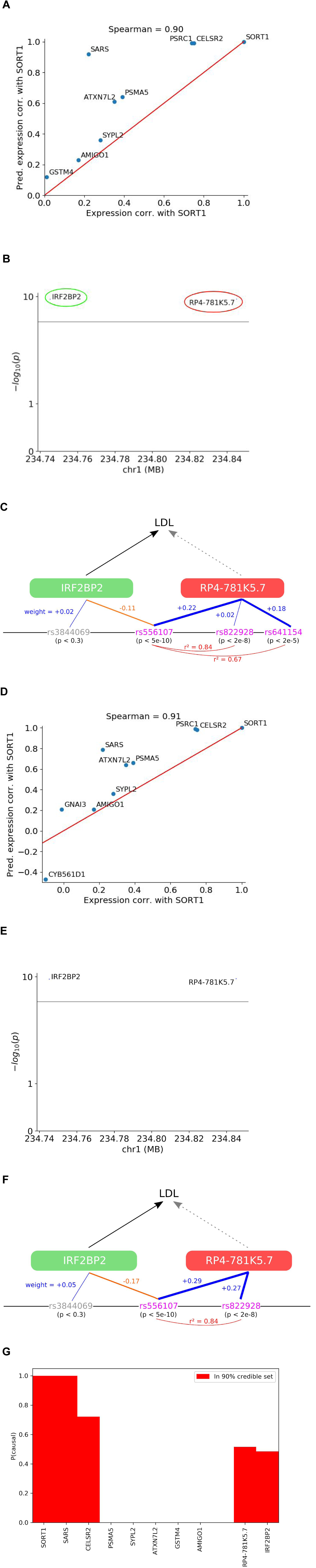
Correlated predicted expression can cause non-causal hits even in the absence of correlated total expression. a) For nearby genes, Fusion predicted expression correlations tend to be higher than total expression correlations, e.g. at the *SORT1* locus. b) Fusion Manhattan plot of the *IRF2BP2* locus, where *RP4-781K5.7* is a likely non-causal hit due to predicted expression correlation with *IRF2BP2*. c) Details of the two genes’ Fusion expression models: a line between a variant’s rs number and a gene indicates the variant is included in the gene’s expression model with either a positive weight (blue) or negative weight (orange), with the thickness of the line increasing with the magnitude of the weight; red arcs indicate LD. Pink rs numbers are GWAS hits (genome-wide-significant or sub-significant) while gray rs numbers are not. d) The equivalent plot to a) for S-PrediXcan. e) The equivalent plot to b) for S-PrediXcan. f) The equivalent plot to c) for S-Predixcan. For clarity, 4 variants with weights less than 0.05 in magnitude for IRF2BP2 (rs2175594, p < 0.02, weight +0.03; rs2439500, p < 0.2, weight = +0.01; rs11588636, p < 0.3, weight = -0.03; rs780256, p < 0.9, weight = -0.03) and 5 variants for RP4-781K5.7 (rs478425, p < 0.01, weight = +0.02; rs633269, p < 0.02, weight = +0.01; rs881070, p < 0.06, weight = -0.02; rs673283, p < 0.1, weight = +0.004; rs9659229, p < 0.1, weight = -0.04) are not shown. g) Estimated causal probability for each significant gene from Fusion at the *SORT1* and *IRF2BP2* loci, according to TWAS gene-based fine-mapping with the FOCUS method.

Predicted expression correlation may lead to non-causal genes being prioritized before causal genes, even if the total expression correlation between the two genes is low (Fig. 5b). This type of confounding has also been observed in the context of gene-set analysis^26^. For instance, *SARS* is the main outlier in Fig. 2b because, despite having a similar Fusion *p*-value to *SORT1*, it has an unexpectedly low total expression correlation of approximately 0.2; yet it is still a strong hit because of its high predicted expression correlation of approximately 0.9 (Fig. 3a); *SARS* is also an outlier in PrediXcan, for the same reason (Fig. 3d). Of course, it is always possible that *SARS* may be an outlier due to having a causal effect of its own.

Another example is the *IRF2BP2* locus in LDL/liver (Fig. 3b). *IRF2BP2* is a gene encoding an inflammation-suppressing regulatory factor with evidence of causality from mouse models (Table 3). *RP4-781K5.7* is a largely uncharacterized long non-coding RNA (lncRNA), and lacks evidence for a causal role in LDL or indeed for having any function at all; most lncRNAs are non-essential for cell fitness^27^ and current evidence is compatible with a model where most non-coding RNAs are non-functional^28^. Nonetheless, it is possible that *RP4-781K5.7* does also have a causal role in LDL. While there is almost no correlation in total expression between the two genes (−0.02), *IRF2BP2*’s Fusion expression model includes a GWAS hit variant, rs556107, with a negative weight while *RP4-781K5.7*’s includes the same variant, as well as two other linked variants, with positive weights (Fig. 3c), resulting in almost perfectly anti-correlated predicted expression between the two genes (−0.94) and significant TWAS associations with LDL for both genes. *IRF2BP2* and *RP4-781K5.7* are also both hits with S-PrediXcan (Fig. 3e) and, as with Fusion, both models put the largest weight on rs556107 but with opposite sign (Fig. 3f).

**Table 3:** Candidate causal genes curated from the literature, with supporting evidence for causality. The strength of evidence for each gene is also stated: strong indicates clear experimental evidence (*SORT1*, *PPARG*, *FADS1-3*, *KPNB1*, *STAT3*), coding loss-of-function or fine-mapped GWAS association (*IRF1*, *CARD9*, *NOD2*) or functional inference (*LPA*) linking the gene to the trait; moderate indicates less direct experimental (*TNKS*) or functional (*ALDH2*, *LPIN3*) evidence, or clear experimental evidence linking the gene to a related trait (*IRF2BP2*); weak indicates disputed evidence of causality, where another gene at the locus is a stronger candidate (*SLC22A4/5*).

| Gene | Trait | Evidence | Details |
| --- | --- | --- | --- |
| *SORT1* | LDL | Strong | In mouse models, overexpression of *SORT1* in liver reduced plasma LDL levels and siRNA knockdown increased plasma LDL levels<sup>23,30</sup>, though in other studies deletion of *SORT1* counter-intuitively reduced, rather than increased, atherosclerosis in mice without affecting plasma LDL levels<sup>31,32,33</sup>. |
| *IRF2BP2* | LDL | Moderate | A loss-of-function variant in *IRF2BP2* has been associated with increased susceptibility to CAD<sup>34</sup>. *IRF2BP2* knockout has been shown in mouse models to increase atherosclerosis, albeit via an inflammatory mechanism<sup>34</sup>. |
| *PPARG* | LDL | Strong | *PPARG* activation increases LDL metabolism via induction of *LDLR* and *CYP7A1*<sup>35</sup>; PPAR agonists decrease glycated LDL uptake into macrophages via regulation of lipoprotein lipase<sup>36</sup>. |
| *LPA* | LDL | Strong | *LPA* is a primary constituent of lipoprotein(a), a class of lipoproteins related to LDL. The LDL GWAS used in this study is a meta-analysis of 60 studies, most of which do not measure LDL levels directly but instead calculate them indirectly using the Friedewald formula, which does not distinguish between LDL and lipoprotein(a) and instead reports the sum of LDL and lipoprotein(a) levels<sup>37</sup>. Thus, although *LPA* abundance may not causally influence true LDL levels, it does causally determine LDL levels as calculated by the Friedewald formula. |
| *TNKS* | LDL | Moderate | Inhibition of *TNKS* inhibits Wnt signalling<sup>38</sup> and upregulates genes involved in cholesterol biosynthesis<sup>39</sup>. Wnt signalling has independently been implicated in lipid homeostasis<sup>40,41</sup>. |
| *FADS1-3* | LDL | Strong | *FADS1*-knockout mice had lower triglyceride and total cholesterol levels<sup>42</sup>. *FADS2*-knockout mice had roughly doubled cholesterol synthesis rate in macrophages<sup>43</sup> and altered levels of multiple cholesterol esters in liver<sup>44</sup>. *FADS3* is least well-characterized but has 52% and 62% sequence homology with *FADS1* and *FADS2*, respectively<sup>45</sup>. |
| *ALDH2* | LDL | Moderate | *ALDH2* is required for alcohol metabolism, and alcohol consumption has long been known to have wide-ranging influences on lipid levels<sup>46</sup>. Both *ALDH2* and another alcohol metabolic enzyme, *ADH1B*, have been associated with alcohol consumption<sup>47</sup>, variants at both loci have been associated with LDL among alcoholic men<sup>48</sup>, and Mendelian randomization using variants near *ADH1B* and other alcohol metabolic enzymes recapitulated the causal role of alcohol consumption on LDL levels<sup>49</sup>, suggesting that *ALDH2* causally influences LDL levels via its effect on alcohol consumption. |
| *KPNB1* | LDL | Strong | *KPNB1* knockdown reduced cellular internalization of fluorescence-labeled LDL<sup>50</sup>. |
| *LPIN3* | LDL | Moderate | *LPIN3* is one of three lipin genes; lipin genes catalyze the synthesis of diacylglycerols<sup>51</sup>, constituents of LDL and other lipoproteins<sup>52</sup> and intermediates in the synthesis of multiple classes of lipids<sup>53</sup>. |
| *SLC22A4/5* | Crohn's | Weak | Also known as *OCTN1/2*, these proteins transport substrates such as ergothioneine and acetylcholine, and a Crohn's-associated variant, L503F, increases *SLC22A4* 's transport efficiency<sup>54,55</sup>. However, the link between altered transport efficiency and disease is unclear, and *IRF1* is a stronger candidate at the locus (see below). |
| *IRF1* | Crohn's | Strong | Genome-wide, variants that increase binding of *IRF1* (a transcriptional activator of the innate immune response) tend to increase Crohn's risk, and vice versa<sup>56</sup>. High-density genotyping of the *IRF1/SLC22A4/5* locus indicates that *IRF1*, but not *SLC22A4/5*, associates with Crohn's disease risk, and *IRF1* expression, but not *SLC22A4* expression, in GI biopsies was increased among Crohn's cases<sup>57</sup>. |
| *CARD9* | Crohn's | Strong | *CARD9* plays critical roles in the innate immune response and has been implicated in a variety of autoimmune conditions<sup>58</sup>; a loss-of-function splice variant in *CARD9* is strongly protective against Crohn's disease<sup>59</sup>. |
| *NOD2* | Crohn's | Strong | Multiple coding variants in *NOD2* are independently associated with Crohn's disease<sup>59,60,61</sup>. |
| *STAT3* | Crohn's | Strong | *STAT3*-knockout mice develop Crohn's-like symptoms<sup>62</sup>. |

To illustrate this phenomenon, we simulated expression and trait data (N_trait_ = 10,000 individuals, N_expression_ = 489 individuals from 1000 Genomes of European ancestry) for 1000 random genomic loci using the FOCUS simulation framework^25^ (see “Suggested best practices and future opportunities”), and conducted TWAS using L2-penalized linear regression (see Methods). As expected, larger predicted expression correlation increased the probability of having a larger TWAS z-score than the causal gene (Table 2). However, this probability remained high even when predicted expression correlation was low, suggesting that predicted expression, though better than true expression, still imperfectly captures co-regulation in the context of TWAS (see next section).

## Shared GWAS variants between gene expression models may lead to spurious TWAS hit genes, even without correlated predicted expression

More generally, pairs of genes may share GWAS variants in their models (or at least share LD partners, where a variant in one gene’s model is in LD with a variant in the other gene’s model) even if they have low predicted expression correlation, since other variants that are distinct between the models may “dilute” the correlation (Fig. 5c). For instance, at the *NOD2* locus for Crohn’s/whole blood, *NOD2* is a known causal gene (Table 3), but 4 other genes are also Fusion TWAS hits (Fig. 4a), none with strong evidence of causality (though rare variants in one gene, *ADCY7*, have been associated with the closely related disease ulcerative colitis but not Crohn’s_29_). The Fusion model for the strongest hit at the locus, *BRD7*, puts most of its weight on rs1872691, which is also the strongest GWAS variant in *NOD2*’s model (Fig. 4b). However, the *NOD2* model puts most of its weight on two other variants, rs7202124 and rs1981760, which are slightly weaker GWAS hits. The result is that even though *BRD7* appears to be a non-causal hit because of co-regulation with *NOD2* (though it is certainly possible that *BRD7* may be causal itself), the overall predicted expression correlation between the two genes is very low (−0.03), as is the total expression correlation (0.05). The same 5 genes are also hits with rediXcan (Fig. 4c), and *NOD2* and *BRD7*’s models share the same rs1872691 variant, just like with Fusion (Fig. 4d).

**Figure 4:**
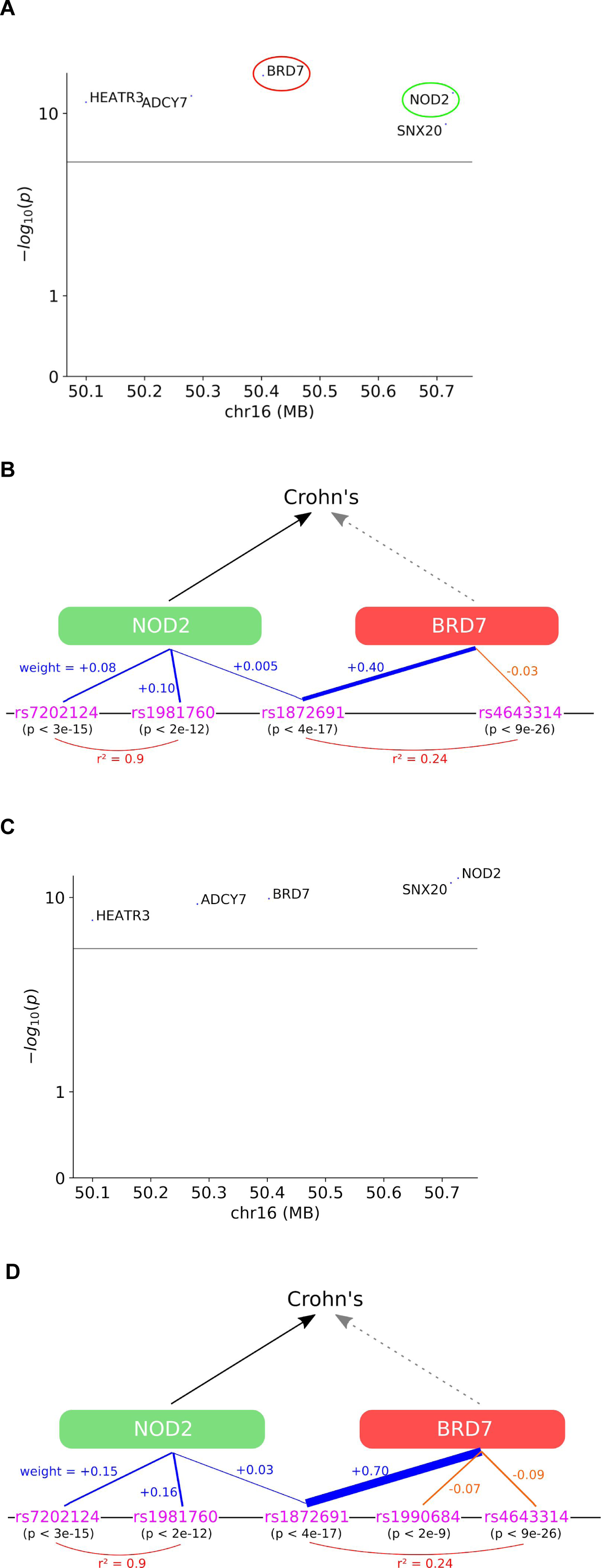
Sharing of GWAS variants between expression models can contribute to non-causal hits even without correlated predicted expression. a) TWAS Manhattan plot of the *NOD2* locus. b) Details of the expression models of *NOD2* and *BRD7*: as in Fig. 3, a line between a variant’s rs number and a gene indicates the variant is included in the gene’s expression model with either a positive weight (blue) or negative weight (orange), with the thickness of the line increasing with the magnitude of the weight; red arcs indicate LD. c) The equivalent plot to a) for S-Predixcan. d) The equivalent plot to b) for S-PrediXcan. For clarity, 5 variants for BRD7 (rs12925755, p < 6e-34, weight = 0.002; rs2066852, p < 3e-10, weight = -0.02; rs17227589, p < 2e-7, weight = -0.02; rs11642187, p < 0.04, weight = +0.007; rs2241258, p < 0.3, weight = -0.05) are not shown.

In the most general case, models need not even share the same GWAS variants (or variants in LD) for there to be spurious non-causal hits (Fig. 5d). For instance, rs4643314, the stronger GWAS hit out of the two variants in *BRD7*’s Fusion model, is neither shared nor in strong LD with any of the variants in *NOD2*’s model, though it is in weak LD with rs1872691 (Fig. 4b). Under the assumption that *NOD2* is the only causal gene at the locus, this suggests that this variant exerts its GWAS effect via *NOD2* and also happens to co-regulate *BRD7*, but that the *NOD2* expression model incorrectly fails to include it (a false negative). One trivial reason for false negatives is variants outside the 500 kb or 1 MB window around the TSS included in the expression modeling (not an issue in this example because rs4643314 is within 500 kb of the *NOD2* TSS), which can be solved by increasing the window size. More problematically, bias in the expression panel could also lead to false negatives (see Discussion). The scenario of multiple distinct GWAS hits might occur even without any false negatives in the expression modeling, e.g. if a variant in *BRD7*’s model (or a variant in LD with it) deleteriously affected the coding sequence of *NOD2* as well as regulating *BRD7*. This bias due to coding variation may occur even despite TWAS not being designed to detect associations mediated by coding variants.

**Figure 5:**
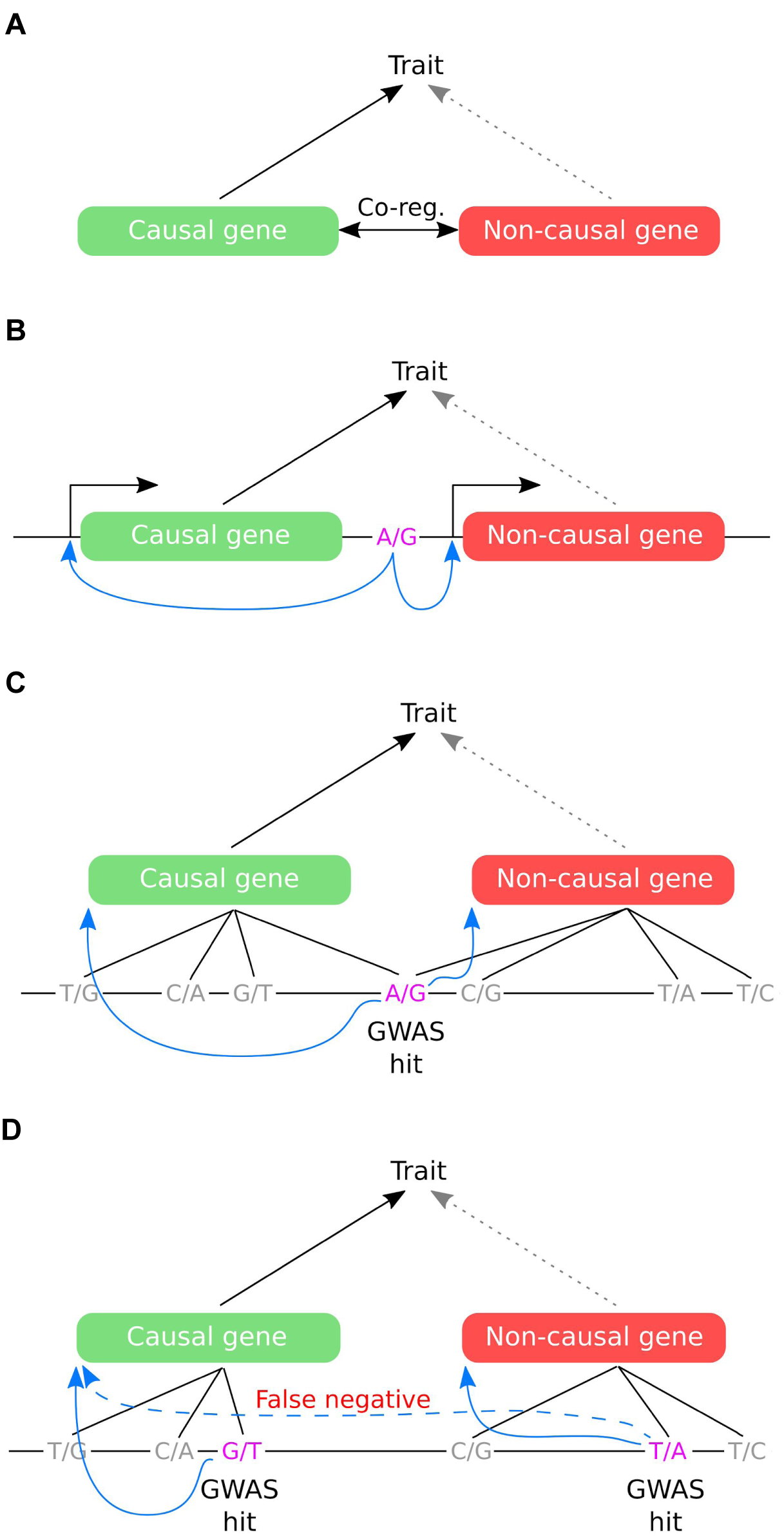
Co-regulation scenarios in TWAS that may lead to non-causal hits, from least to most general. a) Correlated expression across individuals: the causal gene has correlated total expression with another gene, which may become a non-causal TWAS hit. b) Correlated predicted expression across individuals: even if total expression correlation is low, predicted expression correlation may be high if the same variants (or variants in LD) regulate both genes and are included in both models. c) Sharing of GWAS hits: even if the two genes’ models include largely distinct variants and predicted expression correlation is low, only a single shared GWAS hit variant (or variant in LD) is necessary for both genes to be TWAS hits. d) Both models include distinct GWAS hits: in the most general case, the GWAS hits driving the signal at the two genes may not be in LD with each other, for instance if the non-causal gene’s GWAS hit happens to regulate the causal gene as well but this connection is missed by the expression modeling (a false negative), or if the causal gene’s GWAS hit acts via a coding mechanism (not shown).

For methods using GWAS summary statistics (e.g. Fusion and S-PrediXcan), false negatives may also occur due to a mismatch in LD between the expression panel and the GWAS: for instance, a causal gene’s TWAS model for may rely on a variant that is tightly linked to the causal variant in the expression panel, but if this variant is not also linked in the GWAS, the gene may erroneously fail to be detected as a TWAS hit. Conversely, a non-causal gene’s TWAS model may rely on a variant that is linked, in the GWAS but not the expression panel, to a causal variant for a different gene, leading to the non-causal gene being a TWAS hit.

## Using reference gene expression panels from tissues that are less related to the trait introduces bias in TWAS

It is common practice to use reference expression panels in tissues with the largest number of individuals available (e.g. whole blood, lymphoblastoid cell lines) with the goal of maximizing power, even if they are less mechanistically related to the trait. So far, our TWAS case studies have used expression from tissues with a clear mechanistic relationship to the trait: liver for LDL and whole blood for Crohn’s. What if we swap these tissues and use liver for Crohn’s and whole blood for LDL, so that we are using tissues without a clear mechanistic relationship? It is well-known that the architecture of eQTLs differs substantially across tissues: even among strong eQTLs in GTEx (p ~ 1 × 10^−10^), one quarter switch which gene they are most significantly associated with across tissues^18^.

We manually curated candidate causal genes from the literature (Table 3) at 9 LDL/liver and 4 Crohn’s/whole blood multi-hit Fusion TWAS loci and looked at how their hit strengths changed when swapping tissues (Fig. 6). Strikingly, almost every candidate causal gene (9 of 11 for LDL and 5 of 6 for Crohn’s) was no longer a hit in the “opposite” tissue, either because they were not sufficiently expressed (*N* = 4: *PPARG*, *LPA*, *LPIN3*, *SLC22A4*) or because they did not have sufficiently heritable *cis* expression, according to a likelihood ratio test, to be tested by Fusion (*N* = 10: *SORT1*, *IRF2BP2*, *TNKS*, *FADS3*, *ALDH2*, *KPNB1*, *SLC22A5*, *IRF1*, *CARD9*, *STAT3*). This general trend also holds globally, albeit less strongly: genome-wide, 3085 of 5858 LDL/liver genes (53%) drop out when switching to whole blood, and 1202 of 2118 Crohn’s/whole blood genes (57%) drop out when switching to liver. It is important to note that just because a gene does *not* drop out, and is present in both tissues due to shared cross-tissue regulatory architecture, does not necessarily make it more likely to be causal.

**Figure 6:**
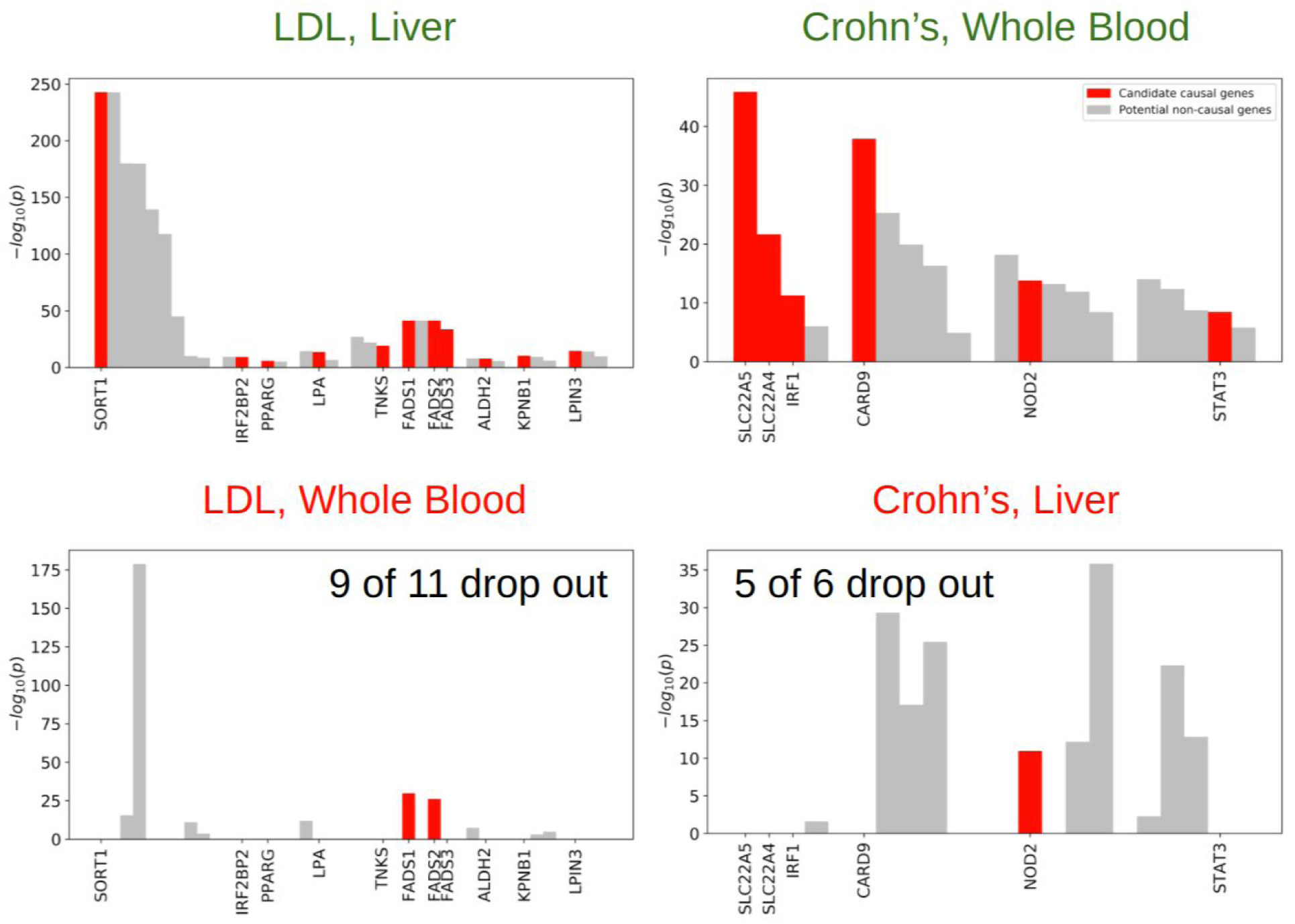
Most candidate causal genes drop out when switching to a tissue with a less clear mechanistic relationship to the trait, due to lack of sufficient expression or sufficiently heritable expression. Fusion TWAS *p*-values at 9 LDL/liver and 4 Crohn’s/whole blood multi-hit loci, when using expression from tissues with a clear (top row) and less clear or absent (bottom row) mechanistic relationship to the trait. Candidate causal genes are labeled and colored in red.

More problematically, 15 other genes at the same loci were still hits (8 in LDL/whole blood and 7 in Crohn’s/liver), and 5 were even strong hits with p < 1 × 10^−20^. This suggests that the strategy of conducting TWAS in a tissue that is sub-optimal for the trait being examined, just because that tissue happens to have a large expression reference panel, is especially problematic because many hit loci may contain only non-causal genes and the causal gene may not even be included in the list of hits.

## TWAS offers improved causal gene prioritization relative to simple baselines

We investigated the performance of TWAS at ranking (prioritizing) causal genes at each of the TWAS hit genes from the previous section. We compared Fusion TWAS to two simple baselines (Table 4): the proximity of each gene’s TSS to the lead (most significant) GWAS variant within 2.5 MB of any gene at the locus (“proximity”), and the median expression across GTEx individuals of each gene in the more mechanistically related tissue, i.e. liver for LDL genes and whole blood for Crohn’s genes (“expression”). By convention, higher rankings are given to genes with greater probabilities of causality (more significant TWAS p-values, closer to the lead GWAS variant, or higher expression). All three methods perform better than a random ranking of the genes at the locus: the mean rank of the 17 candidate causal genes is 3.9, but their mean rank by TWAS is 2.0, by proximity 2.2, and by expression 2.9. Hence, in this simple test, Fusion performs better than both baselines, albeit only slightly better than ranking genes by proximity to the lead GWAS variant.

**Table 4:** Performance comparison of Fusion TWAS, expression and proximity to lead variant at ranking candidate causal genes.

| Candidate causal gene | Number of TWAS hit genes at locus | Rank - TWAS | Rank - proximity | Rank - expression |
| --- | --- | --- | --- | --- |
| *SORT1* | 9 | 1 | 4 | 4 |
| *IRF2BP2* | 2 | 2 | 2 | 2 |
| *PPARG* | 2 | 1 | 1 | 1 |
| *LPA* | 3 | 2 | 3 | 1 |
| *TNKS* | 3 | 3 | 3 | 3 |
| *FADS1* | 4 | 1 | 1 | 2 |
| *FADS2* | 4 | 3 | 2 | 4 |
| *FADS3* | 4 | 4 | 4 | 3 |
| *ALDH2* | 3 | 2 | 1 | 3 |
| *KPNB1* | 3 | 1 | 2 | 3 |
| *LPIN3* | 3 | 1 | 3 | 3 |
| *SLC22A4* | 4 | 2 | 3 | 3 |
| *SLC22A5* | 4 | 1 | 2 | 2 |
| *IRF1* | 4 | 3 | 1 | 4 |
| *CARD9* | 5 | 1 | 1 | 3 |
| *NOD2* | 5 | 2 | 1 | 4 |
| *STAT3* | 5 | 4 | 4 | 5 |
| **Mean** | **3.9** | **2.0** | **2.2** | **2.9** |

## Suggested best practices and future opportunities

We have highlighted two vulnerabilities, co-regulation and tissue bias, that affect the performance of TWAS for causal gene prioritization. In this section, we discuss current best practices and future opportunities for mitigating these vulnerabilities.

One emerging approach to address co-regulation is to repurpose GWAS methods for variant fine-mapping to TWAS, following the analogy between LD in GWAS and co-regulation in TWAS. FOCUS (Fine-mapping Of CaUsal gene Sets)^25^ directly models the predicted expression correlations among genes at a TWAS locus to assign a posterior probability of causality to each gene, and can therefore correct for co-regulation due to predicted expression correlation (Fig. 5b). At the *SORT1* locus, FOCUS includes *SORT1*, *SARS* and *CELSR2* in the 90% credible set; at the *IRF2BP2* locus, FOCUS includes both *IRF2BP2* and *RP4-781K5.7* (Fig. 3g). We recommend using fine-mapping methods such as FOCUS to improve the interpretability of TWAS for causal gene identification, or at a minimum considering the relative association strengths (p-values and effect sizes) of all genes at the locus when interpreting TWAS results.

Nonetheless, we recommend keeping in mind the following caveats which make TWAS fine-mapping more challenging than GWAS fine-mapping. Predicted expression only imperfectly captures *cis* expression, the component of expression driven by variants near the gene; there are sources of both variance and bias in the expression modeling:

- **Finite-sized reference panel:** The main source of variance is the finite size of the reference panel. Fortunately, this can be mitigated with Bayesian methods that explicitly model error in the expression predictions^63^. This variance will become less of an issue in the future as reference panel sizes increase.
- **Pleiotropy across tissues:** Traits rarely act through a single tissue: different genes may be causal in different tissues, so even using a tissue where most genes are causal may introduce bias for the remaining genes that are causal in a different tissue. Fortunately, estimating causal tissues on a per-locus basis is an active area of research^64^, and these approaches could be integrated into TWAS fine-mapping in the future.
- **Cell-type heterogeneity:** Most existing expression panels are gathered for heterogeneous tissues consisting of multiple distinct cell types and states. Genes may only be causal for a single cell type/state within a tissue: for instance, a study that identified *IRX3* and *IRX5* as causal genes at the *FTO* locus found genotype-expression associations in primary preadipocytes, a minority of adipose cells, but not in whole adipose tissue^65^. There may be substantial cell type heterogeneity within and between samples (e.g. due to the presence of blood and immune cells, or genetically-driven differences in the relative proportions of cell types within a tissue), which can also introduce bias. Fortunately, with the advent of single-cell RNA sequencing, reference panels for individual cell types are beginning to be compiled, most prominently through the Human Cell Atlas^66^.
- **Bias in expression quantification:** The time of day, physiological state (e.g. time since eating or exercise, disease status) or cause of death of contributors to the expression panel may also subtly bias measurements: though such covariates may be corrected for by methods such as probabilistic estimation of expression residuals^67^ (PEER), any residual signal from covariates may be captured by a gene’s expression model if variants in the vicinity of the gene are associated with a covariate. There may be other sources of bias that are more difficult to quantify.

To address tissue bias, we recommend, in general, using expression from only the most mechanistically related tissue available as the primary analysis, even if this tissue does not have samples from as many individuals as other tissues. However, it may be advisable to switch to a slightly less related tissue (e.g. from a different region of the brain) if doing so would substantially increase the sample size; the trade-off between tissue bias and sample size should be evaluated on a case-by-case basis. When the most related tissue is not known *a priori* for a particular trait, a recent approach based on LD score regression^68^ can be used to determine the best available tissue from among multiple reference panels. Methods to deal with pleiotropy across tissues and cell-type heterogeneity, discussed above in the context of fine-mapping, can also help mitigate tissue bias. If no sufficiently large reference panels from closely related tissues are available, we recommend a tissue-agnostic analysis that aggregates information across all available tissues^4,69^, for improved prioritization relative to using a single unrelated tissue.

## Discussion

In our case studies, we have generally assumed that the single gene with substantial evidence of causality is the sole causal gene at the locus, with some exceptions of loci with multiple causal candidates of varying degrees of evidence (e.g. *FADS1-3*, *SLC22A4*/*5*/*IRF1*). While this is the most parsimonious explanation, it is possible that other loci also harbor multiple causal genes. Indeed, under an omnigenic model of complex traits^70^, every gene may be causal to some degree, though it is still problematic if TWAS identifies marginally causal genes as strong hits due to co-regulation (effect size inflation). Furthermore, the expression of other genes at the locus may causally contribute to the expression of the causal gene, merely by being actively transcribed, even if the gene is non-coding or its protein product has no causal role^71^.

The vulnerabilities we have explored in TWAS, co-regulation and tissue bias, also apply to other methods that integrate GWAS and expression data, although a thorough exploration of these other methods is beyond the scope of this Perspective. Gene-trait association testing based on Mendelian Randomization^5,6,7^ is vulnerable to non-causal hits because co-regulation, as a form of pleiotropy, violates one of the core assumptions of MR^72^. While the HEIDI test^5^ is designed to correct MR in the case where the two genes have distinct, but linked, causal variants, it does not control for the case where the two genes share the same causal variant. GWAS-eQTL colocalization methods such as Sherlock^8^, coloc^9,10^, QTLMatch^11^, eCaviar^12^, enloc^13^ and RTC^14^ are also vulnerable to this phenomenon. The more tightly a pair of genes is co-regulated in *cis*, the more difficult it becomes to distinguish causality based on GWAS and expression data alone. Our results underscore the need for computational and experimental methods that move beyond using expression variation across individuals to complement TWAS in identifying causal genes at GWAS loci.

## Methods

### TWAS with Fusion

TWAS were performed with the Fusion software using default settings and also including polygenic risk score as a possible model during cross-validation in addition to BLUP, Lasso, and ElasticNet. TWAS p values from Fusion were Bonferroni-corrected according to the number of genes tested in the TWAS when assessing statistical significance. Variants in the STARNET reference panel were filtered for quality control using PLINK73 with the options “--maf 1e-10 --hwe 1e-6 midp --geno”. STARNET expression was processed as described in the STARNET paper^22^, including probabilistic estimation of expression residuals67 (PEER) covariate correction. Because Fusion only supports training on PLINK version 1 hard-call genotype files and not genotype dosages, we trained expression models on only the variants both genotyped in STARNET and either genotyped or imputed in the GWAS, filtering out variants without matching strands between the GWAS and STARNET. Expression models were trained on all remaining variants within 500 kb of a gene’s TSS, using Ensembl v87 TSS annotations for hg1974. Linkage disequilibrium and total and predicted expression correlations were calculated across individuals in STARNET.

### TWAS with S-PrediXcan

To run S-PrediXcan, Elastic Net prediction models and linkage disequilibrium reference were generated using the same PEER-corrected STARNET data from the previous section, filtered to match each GWAS. Variants within 1MB from the TSS and 1MN after the TES were used, to predict genetic features annotated as either protein coding, lincRNA or pseudogene in Ensembl v87.

### Simulations

For the simulations, we sampled independent genomic regions as defined by LDetect75. We then annotated each region with overlapping gene transcription start sites using all available genes in RefSeq v65. To simulate trait and expression panel genotypes, we sampled standardized genotypes using the multivariate normal approximation with mean 0 and covariance defined by the linkage disequilibrium among the 489 individuals estimated from 489 1000 Genomes samples of European ancestry.

Next, we simulated heritable gene expression for all genes at a region with 80% of genes having a single causal eQTL and the remaining 20% having 2 causal eQTLs. Causal eQTLs were preferentially sampled within 50 kb of transcription start sites to exhibit a 50x enrichment on average compared with non-overlapping SNPs. Effect sizes for causal eQTLs were drawn from a normal distribution such that genetic variation explained 20% of variance in total expression.

Finally, given expression at genes causal for the complex trait, we sampled gene-level effect sizes from a normal distribution so that 20% of the variance in trait is explained by gene expression. We randomly assigned one gene at the locus to be causal and looked at what percent of the time other genes at the locus had a larger TWAS z-score than this causal gene, as a function of the predicted expression correlation magnitude with the causal gene.

### Code availability

Code to replicate the post-TWAS analysis is available at https://github.com/Wainberg/TWAS_challenges_and_opportunities. The version of Fusion used for this analysis is available at https://github.com/gusevlab/fusion_twas/tree/9142723485b38610695cea4e7ebb508945ec006c.

### Data availability

GWAS summary statistics are publicly available from the CARDIoGRAMplusC4D consortium and Global Lipids Genetics Consortium. STARNET genotypes are available from Johan LM Björkegren on reasonable request. STARNET expression data is available from dbGAP (accession phs001203.v1.p1).

## Acknowledgements

We gratefully acknowledge Jonathan Pritchard, Hua Tang and members of the Zaitlen lab for helpful discussions. This work was funded in part by the Natural Sciences and Engineering Research Council of Canada (NSERC) (grant PGSD3-476082-2015 to M.W.), Stanford Bio-X Bowes fellowship (to M.W.), Stanford Graduate Fellowship (to N.S.-A.), National Defense Science & Engineering Grant (to N.S.-A.), and NIH grants 1DP2OD022870 and U01HG009431 (to A.K.) and 1U24HG008956 and 5U01HG009080 (to M.A.R.).

## Author Contributions

M.W., M.A.R. and A.K. conceived of the study. M.W., N.M. and A.N.B. performed analyses. N.S.-A., D.A.K. and D.G. provided intellectual input. R.E., A.R., T.Q., K.H. and J.L.M.B. provided assistance with the STARNET dataset. M.A.R. and A.K. supervised the study. M.W., M.A.R. and A.K. wrote the manuscript. All authors reviewed the manuscript.

## Supplementary Information

**Figure S1:**
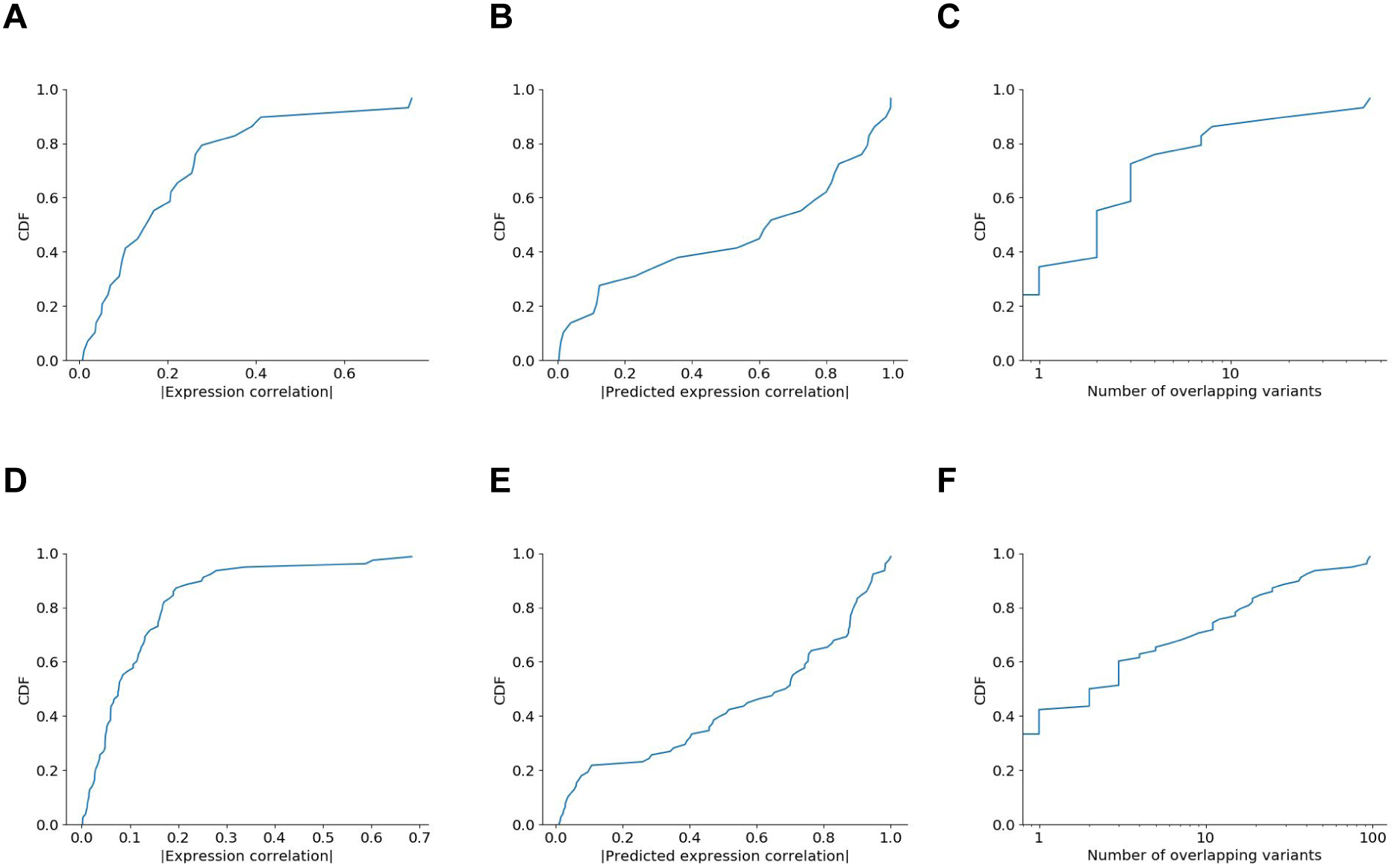
Distributions of co-regulation across putative non-causal genes in multi-hit Fusion TWAS loci. Since many multi-hit loci do not have a clear causal gene or have multiple plausible candidates, we make the approximation that only the most significant gene at each locus is causal. We then plot the cumulative distribution functions (CDFs) of (a, d) expression correlations, (b, e) predicted expression correlations and (c, f) number of shared variants between these most significant genes and all the other genes at their loci, separately for LDL/liver (a-c) and Crohn’s/whole blood (d-f). To collapse these CDFs into a single estimate of the percent of affected non-causal genes (Fig. 1c), we combine genes across the two studies and threshold to correlation r^2^ ≥ 0.2, a threshold commonly used for weak LD in GWAS, or ≥ 1 shared variant. Note that counting only exact sharing of variants does not account for LD, for simplicity.

**Figure S2:**
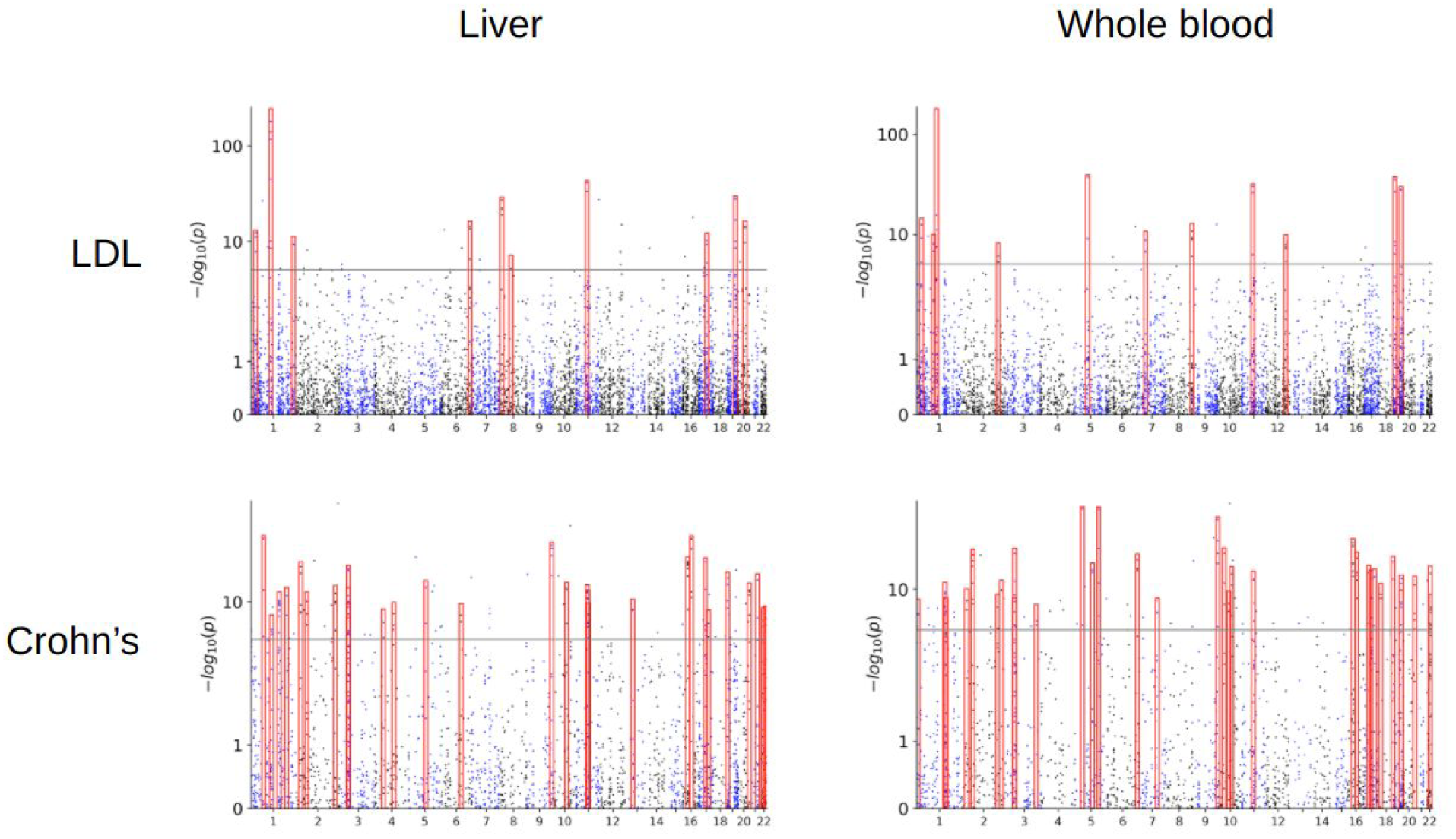
Manhattan plots of the 4 Fusion TWAS conducted in this study. As in Fig. 1, clusters of multiple adjacent TWAS hit genes are highlighted in red.

**Figure S3:**
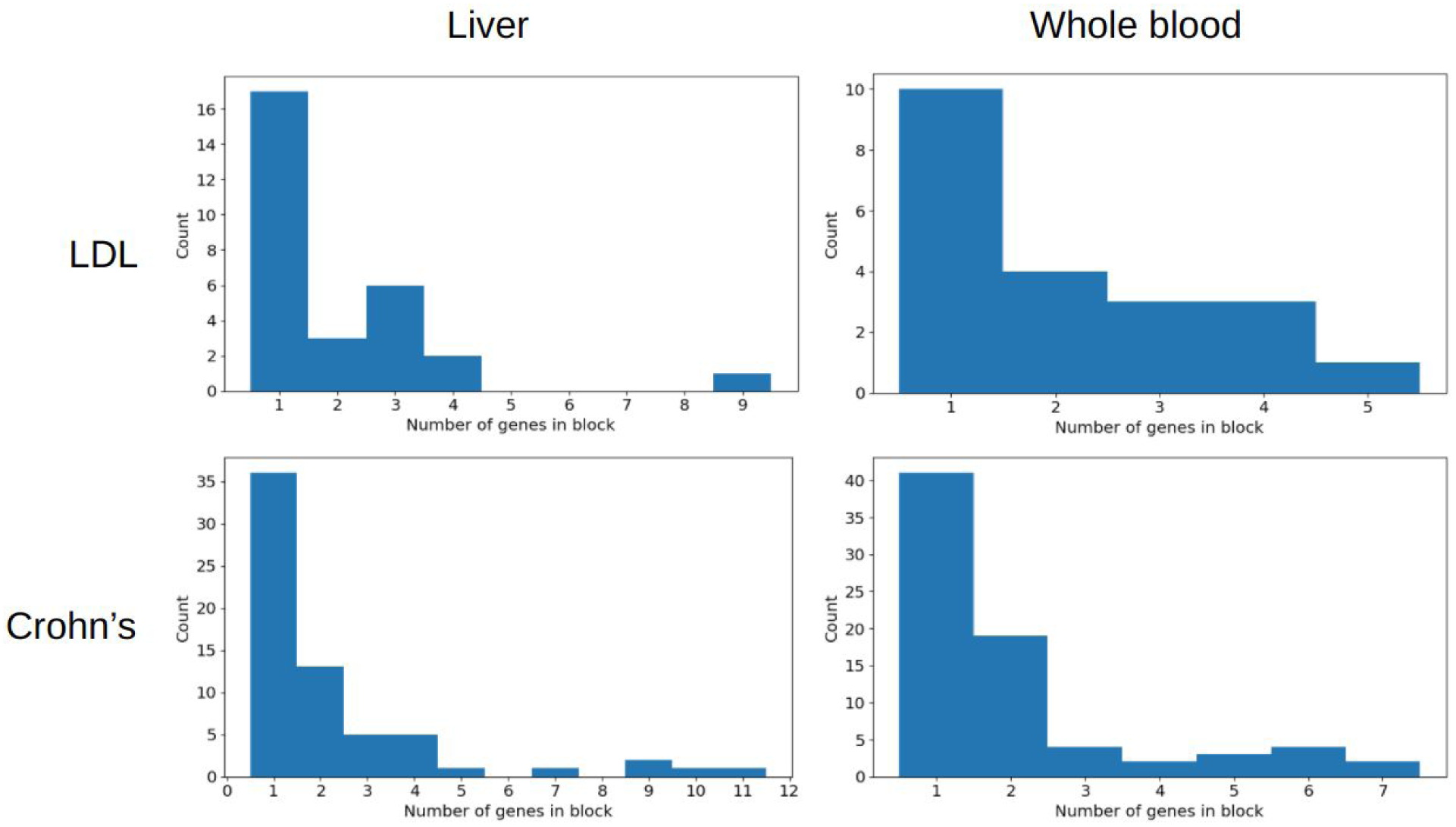
Figure S3: Number of Fusion TWAS hit genes per locus after 2.5-MB clumping.

**Figure S4:**
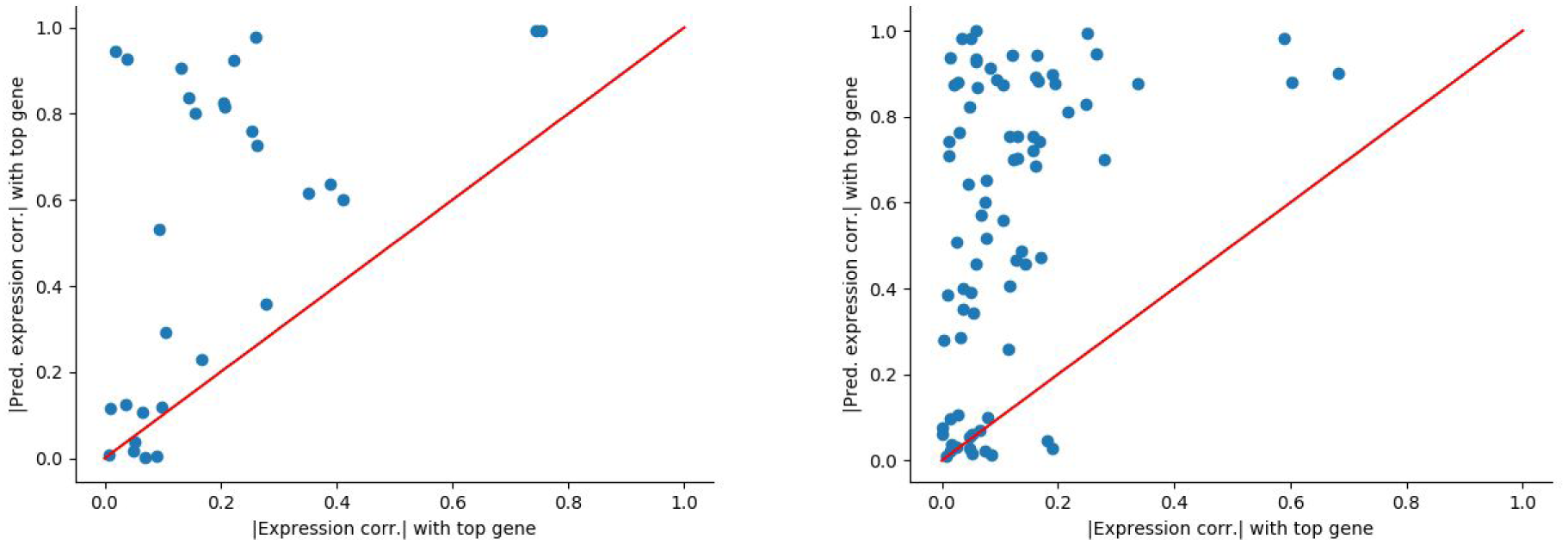
Total versus predicted expression correlation versus the top hit, for all genes in Fusion TWAS multi-hit blocks that are not the top hits. a) Liver, LDL. b) Crohn’s, whole blood. Note that predicted expression correlation is generally higher than total expression correlation, as discussed in the Results section.

